# Activation mapping in multi-center rat sensory-evoked functional MRI datasets using a unified pipeline

**DOI:** 10.1101/2024.09.27.615384

**Authors:** Marie E Galteau, Margaret Broadwater, Yi Chen, Gabriel Desrosiers-Gregoire, Rita Gil, Johannes Kaesser, Eugene Kim, Pervin Kıryağdı, Henriette Lambers, Yanyan Y Liu, Xavier López-Gil, Eilidh MacNicol, Parastoo Mohebkhodaei, Ricardo X N. De Oliveira, Carolina A. Pereira, Henning M Reimann, Alejandro Rivera-Olvera, Erwan Selingue, Nikoloz Sirmpilatze, Sandra Strobelt, Akira Sumiyoshi, Channelle Tham, Raul Tudela, Roël M. Vrooman, Isabel Wank, Yongzhi Zhang, Wessel A van Engelenburg, Jürgen Baudewig, Susann Boretius, Diana Cash, M Mallar Chakravarty, Kai-Hsiang Chuang, Luisa Ciobanu, Gabriel A Devenyi, Cornelius Faber, Andreas Hess, Judith R Homberg, Ileana O Jelescu, Carles Justicia, Ryuta Kawashima, Thoralf Niendorf, Tom WJ Scheenen, Noam Shemesh, Guadalupe Soria, Nick Todd, Lydia Wachsmuth, Xin Yu, Baogui B Zhang, Yen-Yu Ian Shih, Sung-Ho Lee, Joanes Grandjean

**Affiliations:** Donders Institute for Brain, Behaviour, and Cognition, Radboud University, Nijmegen, Radboud University, The Netherlands; Center for Animal MRI, The University of North Carolina at Chapel Hill, 125 Mason Farm Rd., Chapel Hill, The University of North Carolina at Chapel Hill, USA; Neurology, The University of North Carolina at Chapel Hill, 125 Mason Farm Rd., Chapel Hill, The University of North Carolina at Chapel Hill, USA; Biomedical Research Imaging Center, The University of North Carolina at Chapel Hill, 125 Mason Farm Rd., Chapel Hill, The University of North Carolina at Chapel Hill, USA; Translational Neuroimaging and Neural Control Group, High-Field Magnetic Resonance, Max Planck Institute for Biological Cybernetics, 72076, Tuebingen, Max Planck Institute for Biological Cybernetics, Germany; Cerebral Imaging Centre, Douglas Mental Health University Institute, 6875 Boulevard LaSalle, Verdun, Douglas Mental Health University Institute, Canada; Integrated Program in Neuroscience, McGill University, Montreal, McGill University, Canada; Preclinical MRI, Champalimaud Research, Champalimaud Foundation, Avenida de Brasília, Lisbon, Champalimaud Foundation, Portugal; Institute of Experimental and Clinical Pharmacology and Toxicology, FAU Erlangen-Nürnberg, Fahrstrasse 17, 91054 Erlangen, FAU Erlangen-Nürnberg, Germany; Biomarker Research And Imaging in Neuroscience (BRAIN) Centre, Department of Neuroimaging, King’s College London, Coldharbour Lane, London, King’s College London, UK; Experimental Magnetic Resonance Group, Clinic of Radiology, University of Münster, Albert-Schweitzer-Campus 1, Muenster, University of Münster, Germany; Brainnetome CenterBrainnetome Center, Institute of Automation, Chinese Academy of Sciences, Zhongguancun East Road, Beijing, Institute of Automation, Chinese Academy of Sciences, China; Magnetic Imaging Resonance Core Facility, Institut d’Investigacions Biomèdiques August Pi I Sunyer (IDIBAPS), Rosselló, 149-153, Barcelona, Institut d’Investigacions Biomèdiques August Pi I Sunyer (IDIBAPS), Spain; Berlin Ultrahigh Field Facility (B.U.F.F.), Max-Delbrück Center for Molecular Medicine in the Helmholtz Association, Robert-Rössle-Str. 10, Berlin, Max-Delbrück Center for Molecular Medicine in the Helmholtz Association, Germany; NeuroSpin, CEA Saclay, Paris, CEA Saclay, France; Functional Imaging Laboratory, German Primate Center - Leibniz Institute for Primate Research, Kellnerweg 4, Göttingen, German Primate Center - Leibniz Institute for Primate Research, Germany; Faculty of Biology and Psychology, Georg-August University of Göttingen, Göttingen, Georg-August University of Göttingen, Germany; DFG Research Center for Nanoscale Microscopy and Molecular Physiology of the Brain (CNMPB), Göttingen, DFG Research Center for Nanoscale Microscopy and Molecular Physiology of the Brain (CNMPB), Germany; Sainsbury Wellcome Centre, University College London (UCL), 25 Howland Street, London, University College London (UCL), United Kingdom; Institute of Development, Aging and Cancer, Tohoku University, 4-1 Seiryo-machi, Aoba-ku, Sendai, Tohoku University, Japan; National Institutes for Quantum Science and Technology, 4-9-1 Anagawa, Inage-ku, Chiba, National Institutes for Quantum Science and Technology, Japan; Group of Biomedical Imaging, Consorcio Centro de Investigación Biomédica en Red (CIBER) de Bioingeniería, Biomateriales y Nanomedicina (CIBER-BBN), University of Barcelona, Barcelona, University of Barcelona, Spain; Focused Ultrasound Laboratory, Radiology, Brigham and Women’s Hospital, 221 Longwood Ave, Boston, Brigham and Women’s Hos pital, USA; Biological and Biomedical Engineering, McGill University, 3775, rue University, Montreal, McGill University, Canada; Department of Psychiatry, McGill University, 1033 Pine Avenue West, Montreal, McGill University, Canada; Queensland Brain Institute and Centre for Advanced Imaging, University of Queensland, Building 79, St Lucia, University of Queensland, Australia; Dept of Radiology, Lausanne University Hospital (CHUV), rue du Bugnon 46, Lausanne, Lausanne University Hospital (CHUV), Switzerland; Neuroscience and Experimental Therapeutics, Instituto de Investigaciones Biomédicas de Barcelona (IIBB), Consejo Superior de Investigaciones Científicas (CSIC), Institut d’Investigacions Biomèdiques August Pi i Sunyer (IDIBAPS), Rosselló, 161, Barcelona, Institut d’Investigacions Biomèdiques August Pi i Sunyer (IDIBAPS), Spain; Experimental and Clinical Research Center, A Joint Cooperation Between the Charité Medical Faculty and the Max-Delbrück Center for Molecular Medicine in the Helmholtz Association, Berlin, A Joint Cooperation Between the Charité Medical Faculty and the Max-Delbrück Center for Molecular Medicine in the Helmholtz Association, Germany; Department for Medical Imaging, Radboud University Medical Center, PO Box 9101, Nijmegen, Radboud University Medical Cent er, The Netherlands; Erwin L. Hahn Institute for MR Imaging, University of Duisburg-Essen, Essen, University of Duisburg-Essen, Germany; Brain Connectivity and Neuroimaging Lab, Neurosciences Institute, University of Barcelona, Casanova, 143, Barcelona, University of Barcelona, Spain; Athinoula A. Martinos Center for Biomedical Imaging, Massachusetts General Hospital and Harvard Medical School, MA 02129, Charlestown, Massachusetts General Hospital and Harvard Medical School, USA; Biomedical Engineering, The University of North Carolina at Chapel Hill, 125 Mason Farm Rd., Chapel Hill, The University of North Carolina at Chapel Hill, USA

**Author notes:** Corresponding author: dr. Joanes Grandjean.

**Keywords:** sensory-evoked, fMRI, multicenter study, BOLD, Hemodynamic Response Function

## Abstract

Functional Magnetic Resonance Imaging (fMRI) in rodents is pivotal for understanding the mechanisms underlying Blood Oxygen Level-Dependent (BOLD) signals and phenotyping animal models of disorders, amongst other applications. Despite its growing use, comparing rodent fMRI results across different research sites remains challenging due to variations in experimental protocols. Here, we aggregated and analyzed 22 sensory-evoked rat fMRI datasets from 12 imaging centers, totaling scans from 220 rats, to assess the consistency of results across diverse protocols. We applied a standardized preprocessing pipeline and evaluated the impact of different hemodynamic response function models on group and individual level activity patterns. Our analysis revealed inter-dataset variability attributed to differences in experimental design, anesthesia protocols, and imaging parameters. We identified robust activation clusters in all (22/22) datasets. The comparison between stock human models implemented in software and rat-specific models showed significant variations in the resulting statistical maps. Our findings emphasize the necessity for standardized protocols and collaborative efforts to improve the reproducibility and reliability of rodent fMRI studies. We provide open access to all datasets and analysis code to foster transparency and further research in the field.

## Introduction

Functional Magnetic Resonance Imaging (fMRI) in rodents is pivotal to unraveling the mechanisms underlying regional and network level Blood Oxygen Level-Dependent (BOLD) signals^1–6^, phenotyping animal models of disorders^7–9^, and testing pharmacological compounds^10,11^, among many applications. Rodent fMRI offers complementary advantages relative to human fMRI, including the ability to control environmental exposure, study specific genetic influences^12,13^, and evaluate invasive neuromodulatory effects^14,15^. Because of the existence of analogous functional networks between rodent and human brains and the necessity to address critical gaps between mainstream technologies used in rodent and human brain research^16^, the past decade has witnessed a rapid emergence of rodent fMRI studies^17,18^. Nonetheless, this growing community has employed a variety of physiological management procedures during imaging and implemented a wide range of data acquisition and processing protocols to meet different research needs and hardware specifications. These protocol variations affect the quality, reliability, and comparability of the outcomes, impairing an unbiased evaluation across laboratories.

Comparing rodent fMRI across different research sites poses significant challenges^19^. The variability in experimental parameters during animal preparation, data acquisition, and data processing hinders the interoperability of the methods^20,21^. Centers use different restraining protocols, such as awake or anesthetized imaging, anesthetics, physiological control (e.g., free breathing or mechanical ventilation), diverse (multisensory) equipment and paradigms for stimulus-evoked imaging, and a wide range of field strengths and acquisition parameters/protocols^17^. Previously, we have aggregated large dataset collections to compare and to highlight the outcomes of using different experimental parameters. Initially, we scrutinized mouse and rat task-free paradigms^20,22^. We revealed greater than expected variability in the outcomes among the datasets. We identified a rat task-free protocol that is, on average, 60% more sensitive to detecting biologically plausible networks compared to other protocols. Historically, sensory-evoked rat fMRI applications predated task-free protocols^2–6,17,23,24,24–26^. The method has grown from its early origin and is now run in several laboratories. Each uses different equipment and protocols to stimulate rats and evoke neural activity in the corresponding sensory systems (e.g., visual or somatosensory cortex). This raises the question of how comparable the results across diverse laboratories are.

In this preregistered study, we sought to determine how amenable the methods were to processing using a standardized pipeline and state-of-the-art tools: RABIES^27^ open-source preprocessing pipeline for rodent fMRI and the Python-based toolbox Nilearn^28^ for brain response modeling, and how consistent the results are across different sites and experimental protocols. Specifically, we investigated activity patterns at the group and individual level, and we compared the implementation of different hemodynamic response function models. Our endeavor extends beyond data analysis; we are committed to fostering collaboration and dialogue within the community. With an emphasis on transparency and open science, we have provided unrestricted access to datasets and code.

## Results

We aggregated 22 representative rat sensory-evoked fMRI datasets from 12 imaging centers. There were no restrictions on the strain, sex, age, weight, anesthesia, acquisition system, or imaging sequence. Participating laboratories were instructed to provide 10 functional scans with their corresponding anatomical scans and metadata per dataset. Laboratories could supply more than one dataset if stimuli or acquisition parameters differed. In total, we aggregated 220 scans. We observed notable variations in acquisition parameters between datasets, namely rat strain and handling, imaging methodologies, and experimental designs (Figure 1). There was a sex bias, with 58% males and 42% females. The anesthesia protocols for maintenance and magnetic field strength distributions aligned with the current trend in the field^18^, predominantly using Isoflurane and Medetomidine, and 9.4 T or higher magnetic field strengths. The datasets consisted of sensory stimulation of the forepaw, hindpaw, whiskers, or the eyes. The most common type of stimulation was electrical stimulation of the forepaw, represented by 13/22 datasets collected. Overall, acquisition parameters were eminently heterogeneous. Findings should be interpreted within the context of the characteristics of the present population.

**Figure 1:**
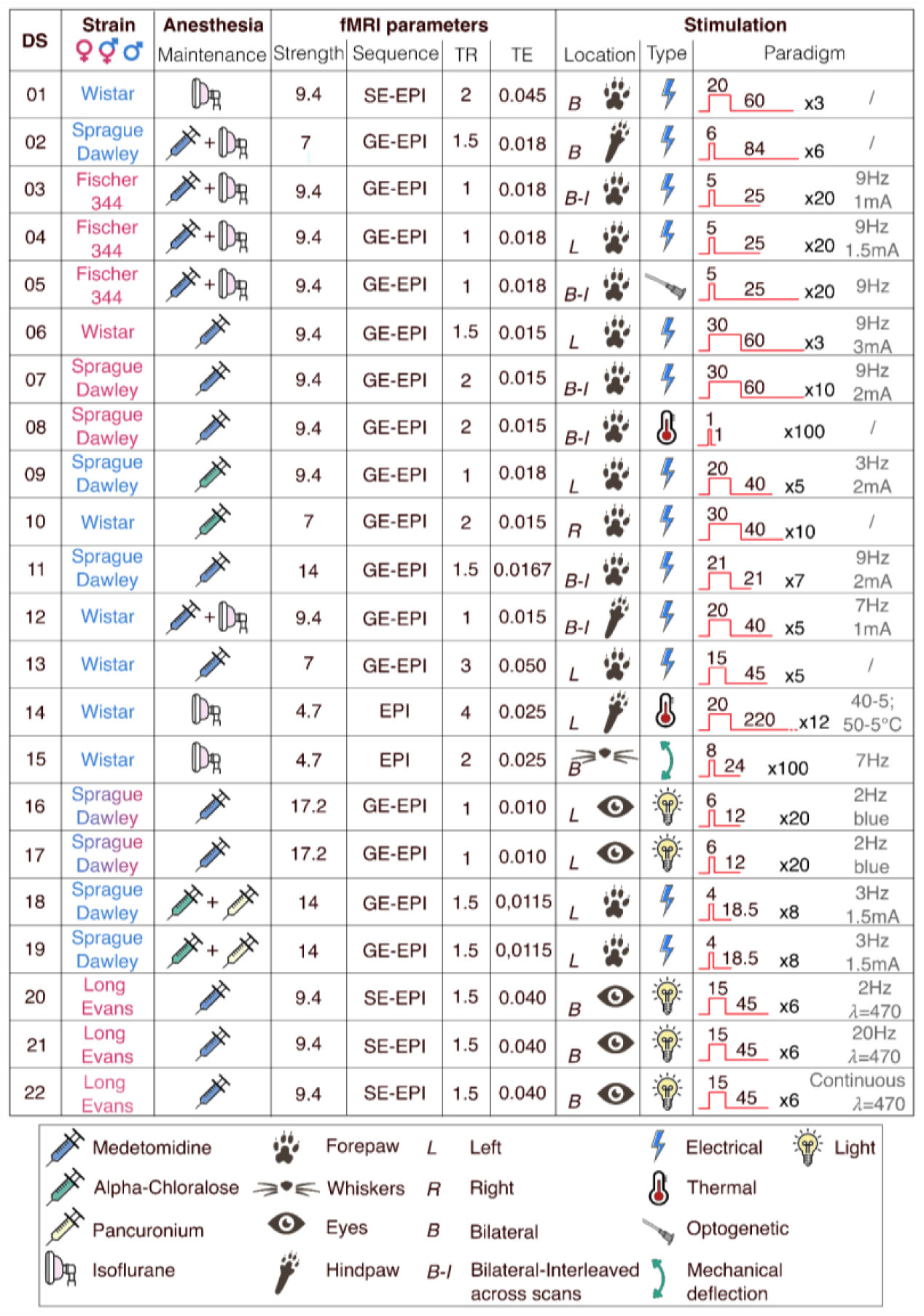
Description of acquisition parameters per dataset (DS). We show the rat strain and sex attributes, maintenance anesthesia type, fMRI parameters such as magnetic field strength in tesla, functional sequence (*SE-EPI: spin-echo echo planar imaging, GE-EPI: gradient-echo echo planar imaging)*, Repetition Time (TR) and Echo Time (TE) in seconds, along with stimulation location, type, and paradigm (in seconds).

We evaluated the consistency of sensory-evoked activity across datasets through a comparative analysis. First, we applied the standardized preprocessing pipeline RABIES^27^, including motion correction, resampling, and registration to the SIGMA rat brain template^29^. We excluded 9/220 subjects due to missing or corrupted functional images and 1/220 for exhibiting failed functional-to-anatomical registration during quality checks after RABIES preprocessing. Following this, we constructed models based on the provided stimulation parameters in Nilearn^28^. We convolved our models using either a rat hemodynamic response function proposed by Lambers et al.^30^ (2-Gammas), a rat hemodynamic response function proposed by Silva et al.^31^ (Peak-span), as well as the default human SPM and Glover response functions implemented within Nilearn. In addition, we examined a Box model based on the block design without convolution. We added six motion parameters and polynomials up to the 3^rd^ degree regressors to the models to account for movement confounds and non-linear low-frequency drifts.

We determined the regions of interest based on the stimulation location: the primary somatosensory cortex forelimb or hindlimb for forepaw or hindpaw stimulations, primary somatosensory cortex barrel field for whisker stimulations, and the superior colliculus for visual stimulations^32,33^. We identified distinct activation clusters within designated regions of interest in all (22/22) group level statistical maps when using the Peak-span rat hemodynamic response function (Figure 2). We noted disparity in cluster intensity and spread. For instance, dataset 13 showed a substantial cluster of activity in the contralateral somatosensory regions with striatal deactivation. In contrast, dataset 06, sharing the same anesthesia, stimulation type, and location, displayed a moderate cluster of activity. To ensure result accuracy and plausibility, we worked with each collaborator to refine the analysis and results. Dataset 22 showed negative activation in the superior colliculus due to the frequency of the visual stimulation (e.g. continuous light), which aligns with the findings of the source center^32^. For datasets acquired with thermal stimulation, namely 08 and 14, we observed diffuse activity patterns consistent with activating the wider pain/saliency matrix^34,35^. Dataset 11 presents negative clusters due to the alignment between the rat model and the actual time series; we found a positive cluster when using the Box model (Figure 4). We also observed activity in the thalamic region for visual stimulation only, while no clearly defined cluster was apparent with other stimulations. From this analysis, we conclude that we can identify activation clusters in all datasets. However, this is accompanied by substantial variability between datasets.

**Figure 2:**
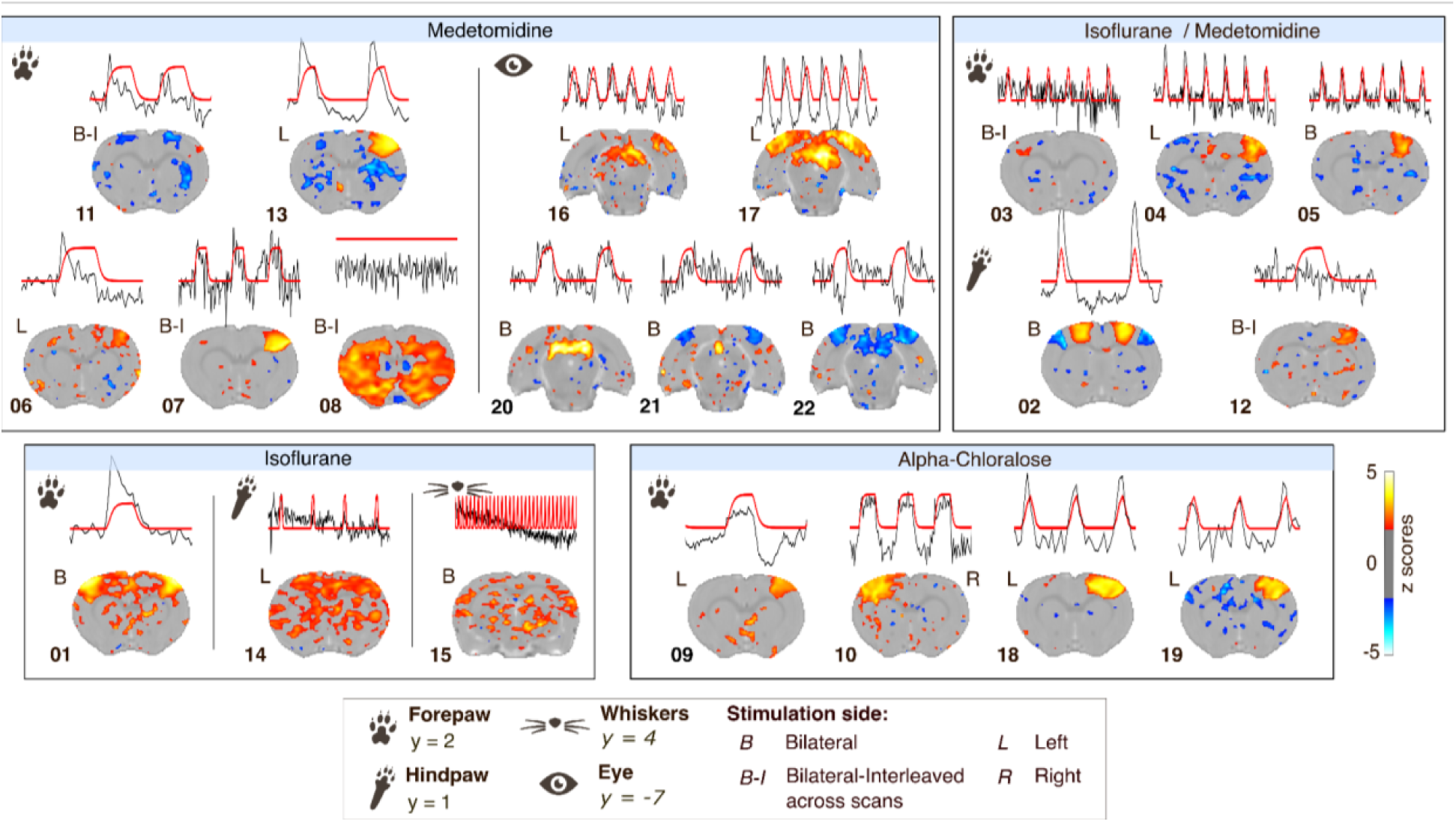
Group level analysis statistical (z-scores, one-sample t-test) maps for datasets processed with the Peak-span hemodynamic response function, accompanied by the modeled response (red) juxtaposed alongside the group-averaged time series from the region of interest (black). The z-scores images are shown as an overlay on the SIGMA template with a threshold set to the significance levels of p < 0.05 uncorrected. The y coordinate along the anterior-posterior axis is given per stimulation location relative to the SIGMA template. Spatial arrangement of maps according to anesthesia and stimulation location.

Analysis is traditionally carried out at the group level. Here, we propose that reliability at the individual level is equally important and can potentially reduce animal use^36^. We investigated the consistency of the individual-evoked response within datasets. In datasets with clear group level clusters, we found that most (but not all) of the individual maps showed robust activation patterns along with 1 or 2 outliers per dataset (e.g., scan with average ROI z-score_Peak-span_ of -0.12 in dataset 01, scan with Z-scorePeak-span = 0.67 in dataset 13, Figure 3). This seems consistent across stimulation methods, anesthesia, and field strengths. We concluded from that analysis that having robust individual level activation is key to high-quality group level maps.

**Figure 3:**
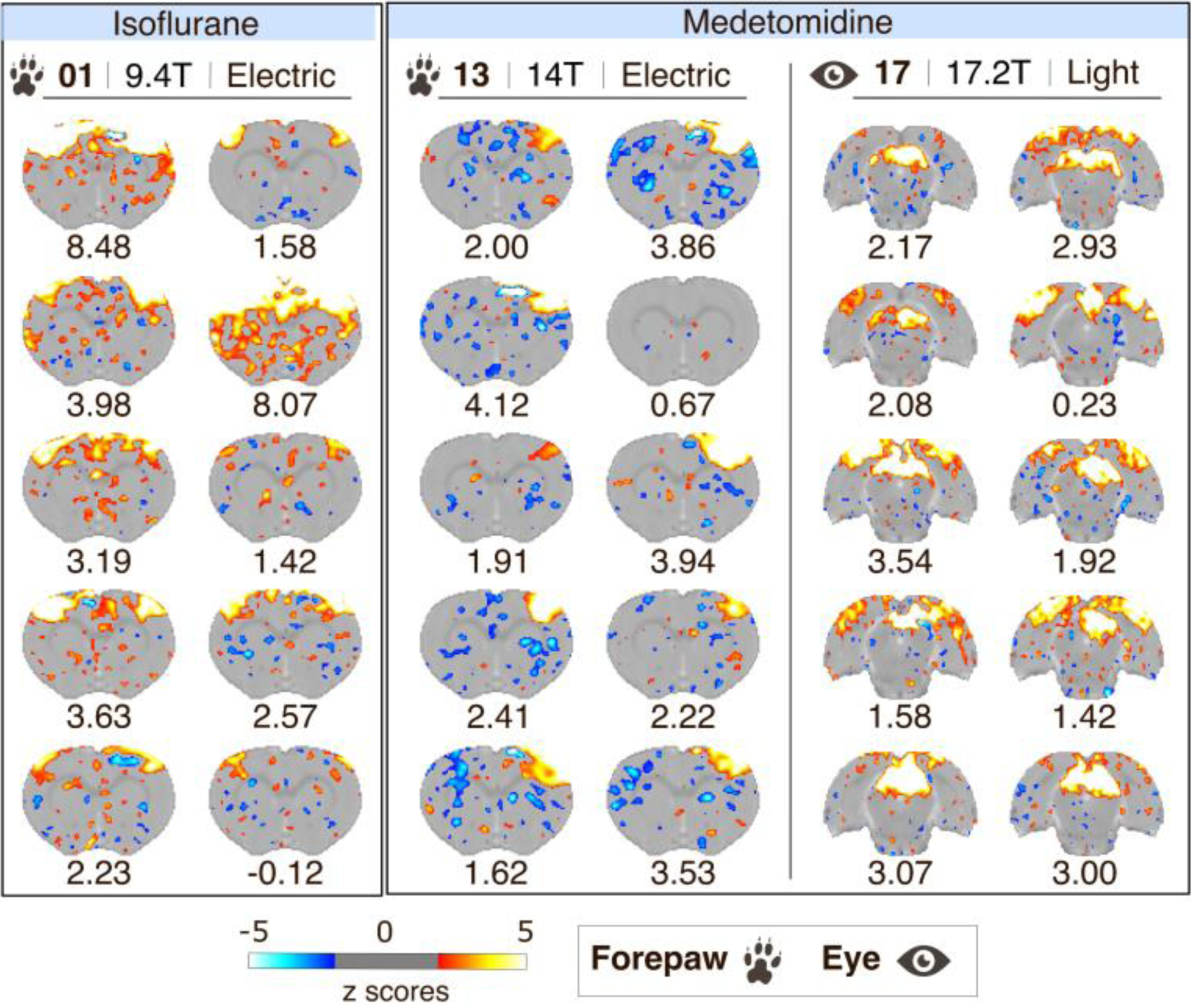
Individual level maps, resulting from the analysis with Peak-span model, for datasets 01, 13 and 17. The spatial arrangement of maps is according to anesthesia, stimulation location, and dataset. The z-scores images are shown as an overlay on the SIGMA template with a threshold set to the significance levels of p < 0.05 uncorrected. The average z-score for the region of interest is provided below the image per scan.

A pivotal aspect of evoked fMRI mapping is the selection of hemodynamic response functions. To date, ad hoc solutions have been tested on datasets from single laboratories^30,31,37,38^. We addressed the impact of hemodynamic response function models by implementing five models into the analysis. These models included the SPM and Glover default human models implemented in Nilearn, a Box model based on the block design, as well as two customized rat models derived from prior research from Silva et al. (Peak-span)^31^ and Lambers et al. (2-Gammas)^30^. The models differed in temporal profiles, peak magnitudes, and rates of decline (Figure 5A). Specifically, the rat models introduced a delay in the peak of activity (Figure 5BC), believed to better fit the BOLD response evoked by sensory stimulation in rats. As a result, we observe variations in statistical maps both at the group level (Figure 4) and individual level when applying different models. The size, amplitude, and polarity of activity clusters changed noticeably when shifting between rat, human and Box models, in a dataset-dependent manner. For instance, in the group level map of dataset 17, the significant positive activity clusters in the primary visual cortex under the rat and Box models (ROI average z_Peak-span_ = 2.20 ± 0.98) became negative when using SPM human models (ROI average z_SPM_ = -1.38 ± 0.68, Figure 4). Other datasets were less impacted by model selection (e.g. dataset 07, ROI average z_2-Gammas_ = 3.75 ± 1.89, z_Peak-span_ = 4.33 ± 1.91, z_Glover_ = 4.70 ± 2.17, z_SPM_ = 4.41 ± 2.03, z_Box_ = 3.62 ± 1.54,). Better fitting models also varied independently of anesthesia or stimulation parameters and appeared instead to be dataset-specific. We found that the Box model provided a better fit, as indicated by higher z-score values, in 12/22 dataset, followed by the Peak-span model (8/22). Surprisingly, human-derived models provided better fits in 1 dataset only (dataset 07). This underlies salient differences brought up by hemodynamic response functions and the need for careful model selection.

**Figure 4:**
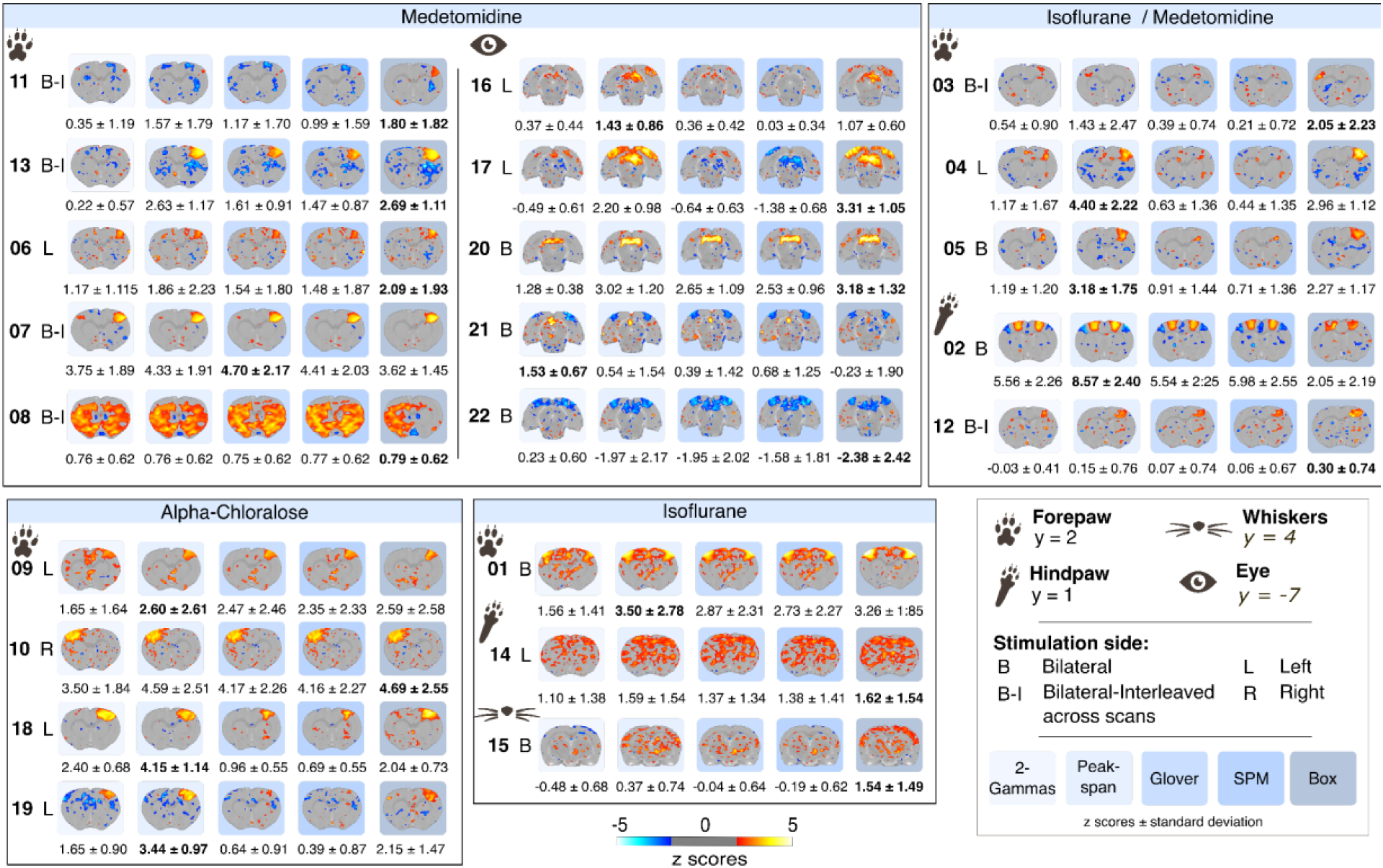
Group level analysis statistical (z-scores, one-sample t-test) maps for datasets processed under each hemodynamic response function, namely, 2-Gammas, Peak-span, Glover, SPM, and Box (from left to right in each row). Thresholds were set to correspond to significance levels of p < 0.05 uncorrected. The y coordinates along the anterior-posterior axis are given per stimulation location relative to the SIGMA template. Spatial arrangement of maps according to anesthesia and stimulation location. The average z-score for the region of interest is provided below the image as the mean ± 1 standard deviation across the scans within the dataset. Bold indicates the highest absolute z-score across models.

**Figure 5:**
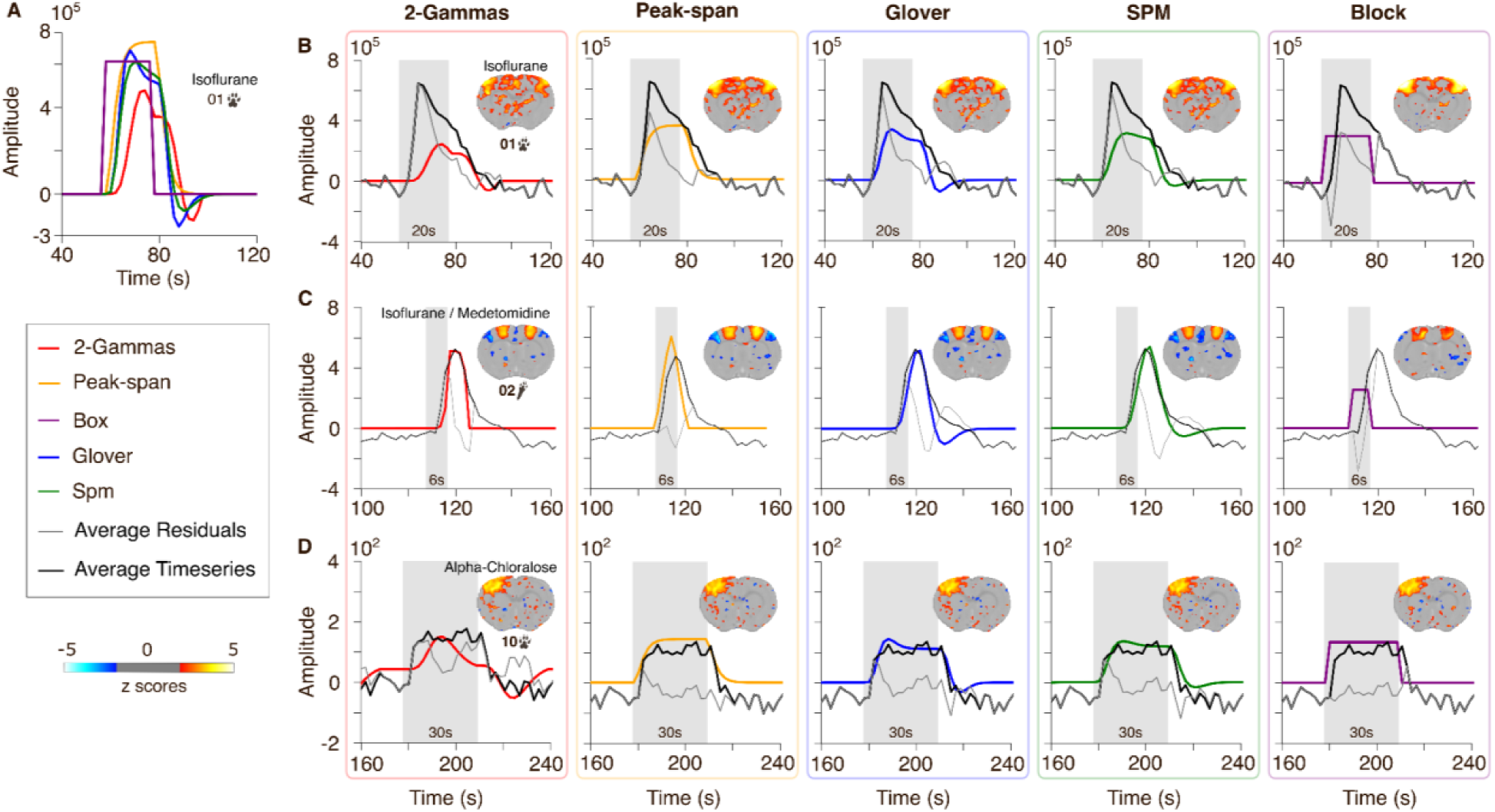
Depiction of the hemodynamic response function models using the stimulation parameters from dataset 01 (A). Namely, the two rat models 2-Gammas and Peak-span, and the two human models Glover and SPM, along with the Box model. Subplots (B-D) show the fit of each model (column) with the time series and the residuals averaged across the 10 animals from dataset 01, 02, and 10 (rows). Shaded area represents the stimulation time.

Next, we examined more closely how the different models fitted the time series. One striking observation from the time series analysis is the presence of an early onset peak in some datasets (datasets 01, 06, 11, 13, Figure 1). This peak was found across anesthesia conditions (isoflurane for 01, medetomidine for the rest), suggesting this may not be due to different neurovascular effects associated with either isoflurane or medetomidine^39^. We examine the model fits in 3 representative datasets: 01, 02, and 10, acquired under isoflurane, isoflurane + medetomidine, and alpha-chloralose, respectively (Figure 5). Dataset 01 presented a noticeable early onset peak, which was not accounted for by any of the models. Among the models, only the non-convolved Box model accounted for the initial fast rise but failed to model the latter phase of the response. Dataset 02 had a response that was best explained by the rat-derived hemodynamic response functions. Finally, dataset 10 was poorly explained by the rat 2-Gammas model but followed more closely by the remaining models, including acceptable fits by the human-derived models. This close examination of the time series further underlies the need for dataset-specific hemodynamic response function selection. To our surprise, the non-convolved Box model appeared a valid heuristic in some instances. We also found that none of the models investigated accounted properly for datasets presenting an early onset rise. Such special cases may need further tailored models for accurate mapping.

## Discussion

There is much to learn from cross-examinations of datasets from multiple centers. Here, we gathered 22 representative datasets, showcasing the diversity in acquisition parameters across centers and datasets, including rat characteristics, animal handling, imaging methodologies, and experimental designs. The first salient observation is that the newly developed preprocessing software, RABIES^27^, could preprocess the vast majority of the data (210/211). This was achieved despite large variations between datasets, including a partial field of view coverage of the brain (e.g. datasets 16 and 17), or inhomogeneous signal distribution. Using the same preprocessing software across studies helps the comparability and reproducibility of the results, and also reduces the need to prepare custom routines per laboratory. The fact that we could preprocess data with large qualitative differences, larger than for task-free applications^20^, further demonstrates the potential of this software. The embedded quality control modules within RABIES also help ensure that datasets processed at different sites undergo comparable scrutiny. Finally, in addition to preprocessing, we implemented two rat hemodynamic response functions within Nilearn. This will help the community seamlessly use these implementations for their future needs.

We applied a standard processing pipeline, which resulted in 22/22 datasets revealing suprathreshold group clusters of activity in the expected regions. This was achieved in a community effort. We carefully reviewed our input parameters and preprocessing steps per dataset with the corresponding data owners. We found that the cluster extents, sign, and amplitude varied between datasets and between HRF models. We found that, most often, the non-convolved Box model yielded better outcomes, followed by the Peak-span model estimated by Silva and colleagues^30^. This was not, however, generalizable across all datasets. Strikingly, we could not identify protocol parameters that would explain the outcome differences. This is due, in part, to the wide range of methods used, regarding both stimulation sites and protocols, but also imaging parameters and equipment. For this reason, we cannot infer the exact causes nor suggest ‘consensus’ protocols. We acknowledge the many parameters that contribute to the enhanced signal, such as anesthesia and physiological maintenance during experimentation^40,41^, but also the selection of stimulation parameters such as the frequency^32^. Our results suggest that the Peak-span rat model is a good heuristic for the analysis of fMRI sensory evoked activity on rodents, especially in the context of electrical stimulation of the forepaw.

Consistency is the key to every scientific endeavor. In this study, we observe that datasets that had consistent activation patterns at the individual level were the ones with more robust activation clusters at the group level. Still, among the more robust datasets, activation was not systematically achieved in all individual scans. We need to better understand the sources creating discrepancies between and within scans. Since individual scans of a dataset are acquired with the same protocols and equipment, we suggest that the variation lies mainly in the physiological parameters^40,42–47^. For instance, the superposition of the spontaneous hemodynamic fluctuation and the evoked response can affect our ability to detect signal changes^48^. This could generate inter-individual variations, including within datasets showcasing evident clusters of activity at the group level. Anesthesia, its impact on physiology and our ability to apply it consistently remains the most likely culprit. Anesthesia protocol comparisons have systematically indicated marked differences in the hemodynamic response amplitude and duration^45,46,49,50^. Finally, within trial habituations may also impact the response amplitude in a sensory modality-specific way and can impact consistency. It is thus credible that our ability to control physiological parameters would yield superior outcomes at the individual level that would be reflected in group level analysis. For this purpose, multimodal approaches, such as joint electrophysiological or calcium recordings, can make a difference in understanding the source of variability in our data. To make a difference here, these methods should also focus on the individual sources of variation. In the meantime, we encourage the implementation and reporting of data quality control, not only related to imaging parameters, to help us collectively improve our ability to detect evoked responses in rodents.

The rat hemodynamic response function exhibits faster temporal kinetics compared to the conventional human hemodynamic response function^31,51,52^. Given these distinctions, using tailored rat hemodynamic response functions for fMRI analysis in rats appears to be a sound heuristic-based decision. Here, we implemented the two rat models, Peak-span^31^ and 2-Gammas^30^, along with a Box model and two human models to allow comparative analysis. However, it is essential to recognize the limitations of the rat models. The Peak-span function model, derived from α-chloralose-anesthetized rats, lacks evidence under different stimulation types or anesthesia conditions. The 2-Gammas function presumes a linear BOLD response, despite demonstrations that the response depends on stimulation time and frequency^30,32^, as well as anesthesia type^53^. Moreover, both functions were derived from somatosensory cortex regions and may be unsuitable for modeling subcortical regions due to hemodynamic deviations from cortical regions^30,54^. Here we find that the assumptions on the model should be more nuanced and do not generalize across datasets.

Interestingly, we found a fast initial peak that was not modeled in any of the HRF functions tested. This fast peak was present in datasets acquired with various anesthesia. The fast nature of the response points towards either neuronal or local vascular components rather than slow modulators such as glial cells that have also been shown to fall off model assumptions^55^. There is substantial evidence within studies that both stimulation frequency and anesthesia duration impact this fast response peak^32,40,56–60^. This underlines the importance of examining datasets from multiple laboratory and stimulation protocols to make sense of this phenomena.

To conclude, we aimed to bring awareness among the research community on the differences and variability between studies and laboratories. A promising starting point to lower the heterogeneity is to build standardized experimental protocols based on successful practices. For now, this heterogeneity underscored the challenge of identifying consistent patterns and limited the generalizability of our findings. For this purpose, we promote collaboration and information sharing among researchers. Researchers should prioritize transparency by including detailed quality assessment measures when reporting results. We also recommend providing open access to the data, to allow scrutiny by peers, facilitate a deeper understanding of the findings, and encourage constructive feedback. The ultimate goal is to record robust and reproducible evoked responses in the rodent brain to accelerate our understanding of the BOLD phenomena, and its downstream mechanisms, but also how this can be used to inform on brain disorders.

## Methods

### Pre-registration

This study was preregistered (https://doi.org/10.17605/OSF.IO/8VY9R). Due to technical limitations, we deviated from the preregistration by not implementing NORDIC correction.

### Data collection

We asked members of the animal MRI community to share datasets through mentions at conferences, social media, and personal invitations. To obtain one or more datasets representative of the acquisitions’ procedures of the source laboratory, we requested datasets including 10 pairs of anatomical and functional scans each, without restrictions on strain, sex, age, weight, anesthesia, acquisition system, or imaging sequence. In total, we gathered 22 datasets representative from 12 imaging centers. We excluded 9 scans due to corrupted or poor raw data.

### Preprocessing

We converted individual datasets to Brain Imaging Data Structure (BIDS) using BrkRaw^61^, a python module to access raw data acquired from Bruker Biospin preclinical MRI scanner, and custom scripts. We ensured that the voxel size and the orientation were specified correctly in the image headers. To account for the T1 effects for acquisitions in the absence of dummy scans, we removed the first; 5 volumes in datasets 03, 04, and 05, 1 volume in dataset 14, and 2 volumes in dataset 15, following the recommendations of the originating laboratories. Further, we flipped the x-axis of 15 individual scans for 6 different datasets with alternating stimulations on the left and right paw, to ensure the activity clusters were located on a consistent side within datasets. To ease image registration, we manually cropped the field of view of 17 individual scans. Subsequently, we preprocessed all scans through RABIES, an open source preprocessing pipeline for rodent fMRI (version: 0.4.8)^27^. The pre-processed steps included motion correction, rigid functional-anatomical registration, non-linear anatomical registration to the SIGMA rat template^29^, and a common space resampling to 0.3 × 0.3 × 0.3 mm^3^, omitting smoothing. No bandpass filter or spatial smoothing was carried out at this stage. To address misregistration instances, we incrementally added autoBox, N4 inhomogeneity correction, and rigid functional-anatomical image alignment options as implemented within RABIES. We carried out rigorous visual quality control checks on the registrations for each scan. Exclusion criteria included raw data with significant artifacts leading to misregistration during preprocessing steps.

### Data analysis

We performed individual and group level analyzes in Nilearn (version: 0.10.0)^28^, using motion parameters as confounds and spatial smoothing with a 0.45 mm^2^ full width at half maximum smoothing kernel. Registered functional scan outputs from RABIES^27^ were processed with Nilearn. We used the design provided by the originating laboratories to construct four models based on different hemodynamic response functions. Namely the two defaults: Glover and SPM without derivatives, a Box model based on the block design without convolution, and two custom rat functions based on previous studies: 2-Gammas and Peak-span. The 2-Gammas rat function aligns with the parameters of the general rat hemodynamic response function, as outlined by Lambers et al.^30^. Similarly, the Peak-span rat function has been defined by a full width at half maximum of 2.18 and a time to peak of 1.92, as described by Silva et al.^31^. We used motion parameters and third-order polynomials as co-regressors to account for motion and drift artifacts. Group level maps were generated using a one-sample t-test on the individual level parameter estimate maps. Time series and parameter estimates were extracted using the SIGMA atlas. individual level and group level maps are represented as z-statistical maps with a threshold set to p < 0.05 uncorrected. The maps are shown as color-coded overlays over the SIGMA template. We used the SIGMA rat template to display statistical maps. We used nilearn *NiftiLabelMasker* function to extract signals from regions of interest (e.g. timeseries, residuals and z-scores). These regions were determined based on the stimulation location: the primary somatosensory cortex forelimb or hindlimb, primary somatosensory cortex barrel field for whisker stimulations, and the superior colliculus for visual stimulations^32,33^. We averaged the residuals and z-scores for the region of interest across subjects within datasets, and we calculated the z-scores standard deviation.

## Data availability

The pre-registration is available under the terms of the CC0 license (https://doi.org/10.17605/OSF.IO/8VY9R). The raw data in BIDS format is available under the terms of the CC0 license (doi:10.18112/openneuro.ds005534.v1.0.0).

## Code availability

The code for this project is available under the terms of the Apache-2 license (https://github.com/grandjeanlab/multirat_se).

## Acknowledgment

This project was kindly supported by the Dutch Research Council (OCENW.KLEIN.334, OSF23.1.037), National Institute of Health (K01EB023983, T32 AA007573), Deutsche Forschungsgemeinschaft (406818964), UK Biotechnology and Biological Sciences Research Council (BBSRC, BB/N009088/1), UK Medical Research Council (MR/N013700/1), European Research Council (ERC; agreement No. PI18/00893, 896245, 679058), Ministerio de Economía y Competitividad (DPI2015-64358-C2-2-R), Fonds de recherche du Québec, Interdisciplinary Center for Clinical Research Münster (PIX), Fundação para a Ciência e Tecnologia (Portugal, project 275-FCT-PTDC/BBB-IMG/5132/2014).

## References

1. Logothetis, N. K. What we can do and what we cannot do with fMRI. Nature 453, 869–878 (2008).

2. Hyder, F. et al. Quantitative functional imaging of the brain: towards mapping neuronal activity by BOLD fMRI. NMR Biomed. 14, 413–431 (2001).

3. Keilholz, S. D., Silva, A. C., Raman, M., Merkle, H. & Koretsky, A. P. Functional MRI of the rodent somatosensory pathway using multislice echo planar imaging. Magn. Reson. Med. 52, 89–99 (2004).

4. Kida, I., Kennan, R. P., Rothman, D. L., Behar, K. L. & Hyder, F. High-resolution CMR(O2) mapping in rat cortex: a multiparametric approach to calibration of BOLD image contrast at 7 Tesla. J. Cereb. Blood Flow Metab. Off. J. Int. Soc. Cereb. Blood Flow Metab. 20, 847–860 (2000).

5. Peeters, R. R., Tindemans, I., De Schutter, E. & Van der Linden, A. Comparing BOLD fMRI signal changes in the awake and anesthetized rat during electrical forepaw stimulation. Magn. Reson. Imaging 19, 821–826 (2001).

6. Duong, T. Q., Silva, A. C., Lee, S. P. & Kim, S. G. Functional MRI of calcium-dependent synaptic activity: cross correlation with CBF and BOLD measurements. Magn. Reson. Med. 43, 383– 392 (2000).

7. Gozzi, A. & Zerbi, V. Modeling Brain Dysconnectivity in Rodents. Biol. Psychiatry 93, 419–429 (2023).

8. Coppola, A. et al. Imaging in experimental models of diabetes. Acta Diabetol. 59, 147–161 (2022).

9. Fang, X. et al. Longitudinal characterization of cerebral hemodynamics in the TgF344-AD rat model of Alzheimer’s disease. GeroScience 45, 1471–1490 (2023).

10. Xi, Z.-X., Wu, G., Stein, E. A. & Li, S.-J. Opiate tolerance by heroin self-administration: an fMRI study in rat. Magn. Reson. Med. 52, 108–114 (2004).

11. Jonckers, E., Van der Linden, A. & Verhoye, M. Functional magnetic resonance imaging in rodents: an unique tool to study *in vivo* pharmacologic neuromodulation. Curr. Opin. Pharmacol. 13, 813–820 (2013).

12. Leong, A. T. L. et al. Optogenetic fMRI interrogation of brain-wide central vestibular pathways. Proc. Natl. Acad. Sci. U. S. A. 116, 10122–10129 (2019).

13. Reinwald, J. R. et al. Psilocybin-induced default mode network hypoconnectivity is blunted in alcohol-dependent rats. Transl. Psychiatry 13, 392 (2023).

14. Ciobanu, L. et al. fMRI contrast at high and ultrahigh magnetic fields: insight from complementary methods. NeuroImage 113, 37–43 (2015).

15. Shih, Y.-Y. I., Yash, T. V., Rogers, B. & Duong, T. Q. FMRI of deep brain stimulation at the rat ventral posteromedial thalamus. Brain Stimulat. 7, 190–193 (2014).

16. Xu, N. et al. Functional Connectivity of the Brain Across Rodents and Humans. Front. Neurosci. 16, 816331 (2022).

17. Mandino, F. et al. Animal Functional Magnetic Resonance Imaging: Trends and Path Toward Standardization. *Front*. Neuroinformatics 13, 78 (2019).

18. Huang, J. et al. The current status and trend of the functional magnetic resonance combined with stimulation in animals. Front. Neurosci. 16, 963175 (2022).

19. Carp, J. The secret lives of experiments: Methods reporting in the fMRI literature. NeuroImage 63, 289–300 (2012).

20. Grandjean, J. et al. A consensus protocol for functional connectivity analysis in the rat brain. Nat. Neurosci. 26, 673–681 (2023).

21. Desrosiers-Gregoire, G., Devenyi, G. A., Grandjean, J. & Chakravarty, M. M. Rodent Automated Bold Improvement of EPI Sequences (RABIES): A Standardized Image Processing and Data Quality Platform for Rodent fMRI. http://biorxiv.org/lookup/doi/10.1101/2022.08.20.504597 (2022) doi:10.1101/2022.08.20.504597.

22. Grandjean, J. et al. Common functional networks in the mouse brain revealed by multi-centre resting-state fMRI analysis. NeuroImage 205, 116278 (2020).

23. Liu, Z. M., Schmidt, K. F., Sicard, K. M. & Duong, T. Q. Imaging oxygen consumption in forepaw somatosensory stimulation in rats under isoflurane anesthesia. Magn. Reson. Med. 52, 277–285 (2004).

24. Lee, S.-P., Silva, A. C. & Kim, S.-G. Comparison of diffusion-weighted high-resolution CBF and spin-echo BOLD fMRI at 9.4 T. Magn. Reson. Med. 47, 736–741 (2002).

25. Silva, A. C. & Koretsky, A. P. Laminar specificity of functional MRI onset times during somatosensory stimulation in rat. Proc. Natl. Acad. Sci. U. S. A. 99, 15182–15187 (2002).

26. Silva, A. C., Lee, S. P., Yang, G., Iadecola, C. & Kim, S. G. Simultaneous blood oxygenation level-dependent and cerebral blood flow functional magnetic resonance imaging during forepaw stimulation in the rat. J. Cereb. Blood Flow Metab. Off. J. Int. Soc. Cereb. Blood Flow Metab. 19, 871–879 (1999).

27. Desrosiers-Grégoire, G., Devenyi, G. A., Grandjean, J. & Chakravarty, M. M. A standardized image processing and data quality platform for rodent fMRI. Nat. Commun. 15, 6708 (2024).

28. Abraham, A. et al. Machine learning for neuroimaging with scikit-learn. *Front*. Neuroinformatics 8, (2014).

29. Barrière, D. A. et al. The SIGMA rat brain templates and atlases for multimodal MRI data analysis and visualization. Nat. Commun. 10, 5699 (2019).

30. Lambers, H. et al. A cortical rat hemodynamic response function for improved detection of BOLD activation under common experimental conditions. NeuroImage 208, 116446 (2020).

31. Silva, A. C., Koretsky, A. P. & Duyn, J. H. Functional MRI impulse response for BOLD and CBV contrast in rat somatosensory cortex. Magn. Reson. Med. 57, 1110–1118 (2007).

32. Gil, R., Valente, M. & Shemesh, N. Rat superior colliculus encodes the transition between static and dynamic vision modes. Nat. Commun. 15, 849 (2024).

33. Dinh, T. N. A., Jung, W. B., Shim, H.-J. & Kim, S.-G. Characteristics of fMRI responses to visual stimulation in anesthetized vs. awake mice. NeuroImage 226, 117542 (2021).

34. Wank, I., Kutsche, L., Kreitz, S., Reeh, P. & Hess, A. Imaging the influence of peripheral TRPV1-signaling on cerebral nociceptive processing applying fMRI-based graph theory in a resiniferatoxin rat model. PLOS ONE 17, e0266669 (2022).

35. Hess, A., Sergejeva, M., Budinsky, L., Zeilhofer, H. U. & Brune, K. Imaging of hyperalgesia in rats by functional MRI. Eur. J. Pain 11, 109–109 (2007).

36. Grandjean, J., Lake, E. M. R., Pagani, M. & Mandino, F. What N Is N-ough for MRI-Based Animal Neuroimaging? eNeuro 11, (2024).

37. Martindale, J. et al. The Hemodynamic Impulse Response to a Single Neural Event. J. Cereb. Blood Flow Metab. 23, 546–555 (2003).

38. Khan, R., Dunn, A. K., Duong, T. Q. & Ress, D. Measurements and Modeling of Transient Blood Flow Perturbations Induced by Brief Somatosensory Stimulation. Open Neuroimaging J. 5, 96–104 (2011).

39. Fukuda, M., Vazquez, A. L., Zong, X. & Kim, S.-G. Effects of the α2-adrenergic receptor agonist dexmedetomidine on neurovascular responses in somatosensory cortex. Eur. J. Neurosci. 37, 80–95 (2013).

40. Sirmpilatze, N., Baudewig, J. & Boretius, S. Temporal stability of fMRI in medetomidine-anesthetized rats. Sci. Rep. 9, 16673 (2019).

41. Bol CJJG, null, Danhof, M., Stanski, D. R. & Mandema, J. W. Pharmacokinetic-pharmacodynamic characterization of the cardiovascular, hypnotic, EEG and ventilatory responses to dexmedetomidine in the rat. J. Pharmacol. Exp. Ther. 283, 1051–1058 (1997).

42. Schroeter, A., Schlegel, F., Seuwen, A., Grandjean, J. & Rudin, M. Specificity of stimulus-evoked fMRI responses in the mouse: The influence of systemic physiological changes associated with innocuous stimulation under four different anesthetics. NeuroImage 94, 372–384 (2014).

43. Pawela, C. P. et al. A protocol for use of medetomidine anesthesia in rats for extended studies using task-induced BOLD contrast and resting-state functional connectivity. NeuroImage 46, 1137–1147 (2009).

44. Weber, R., Ramos-Cabrer, P., Wiedermann, D., van Camp, N. & Hoehn, M. A fully noninvasive and robust experimental protocol for longitudinal fMRI studies in the rat. NeuroImage 29, 1303–1310 (2006).

45. Wei, Z. et al. Toward accurate cerebral blood flow estimation in mice after accounting for anesthesia. Front. Physiol. 14, 1169622 (2023).

46. Le, T. T., Im, G. H., Lee, C. H., Choi, S. H. & Kim, S.-G. Mapping cerebral perfusion in mice under various anesthesia levels using highly sensitive BOLD MRI with transient hypoxia. Sci. Adv. 10, eadm7605 (2024).

47. Cerri, D. H. et al. Distinct neurochemical influences on fMRI response polarity in the striatum. Nat. Commun. 15, 1916 (2024).

48. Saka, M., Berwick, J. & Jones, M. Inter-Trial Variability in Sensory-Evoked Cortical Hemodynamic Responses: The Role of the Magnitude of Pre-Stimulus Fluctuations. Front. Neuroenergetics 4, (2012).

49. Schlegel, F., Schroeter, A. & Rudin, M. The hemodynamic response to somatosensory stimulation in mice depends on the anesthetic used: Implications on analysis of mouse fMRI data. NeuroImage 116, 40–49 (2015).

50. You, T., Im, G. H. & Kim, S.-G. Characterization of brain-wide somatosensory BOLD fMRI in mice under dexmedetomidine/isoflurane and ketamine/xylazine. Sci. Rep. 11, 13110 (2021).

51. de Zwart, J. A. et al. Temporal dynamics of the BOLD fMRI impulse response. NeuroImage 24, 667–677 (2005).

52. Chao, T.-H. H. et al. Computing hemodynamic response functions from concurrent spectral fiber-photometry and fMRI data. Neurophotonics 9, 032205 (2022).

53. Steiner, A. R., Rousseau-Blass, F., Schroeter, A., Hartnack, S. & Bettschart-Wolfensberger, R. Systematic Review: Anesthetic Protocols and Management as Confounders in Rodent Blood Oxygen Level Dependent Functional Magnetic Resonance Imaging (BOLD fMRI)-Part B: Effects of Anesthetic Agents, Doses and Timing. Anim. Open Access J. MDPI 11, 199 (2021).

54. Pawela, C. P. et al. Modeling of region-specific fMRI BOLD neurovascular response functions in rat brain reveals residual differences that correlate with the differences in regional evoked potentials. NeuroImage 41, 525–534 (2008).

55. Schulz, K. et al. Simultaneous BOLD fMRI and fiber-optic calcium recording in rat neocortex. Nat. Methods 9, 597–602 (2012).

56. Kim, T., Masamoto, K., Fukuda, M., Vazquez, A. & Kim, S.-G. Frequency-dependent Neural Activity, CBF, and BOLD fMRI to Somatosensory Stimuli in Isoflurane-anesthetized Rats. NeuroImage 52, 224–233 (2010).

57. Sanganahalli, B. G., Herman, P. & Hyder, F. Frequency-dependent tactile responses in rat brain by fMRI. NMR Biomed. 21, 410–416 (2008).

58. Masamoto, K., Kim, T., Fukuda, M., Wang, P. & Kim, S.-G. Relationship between Neural, Vascular, and BOLD Signals in Isoflurane-Anesthetized Rat Somatosensory Cortex. Cereb. Cortex 17, 942–950 (2006).

59. Lai, H.-Y., Albaugh, D. L., Kao, Y.-C. J., Younce, J. R. & Shih, Y.-Y. I. Robust deep brain stimulation functional MRI procedures in rats and mice using an MR-compatible tungsten microwire electrode. Magn. Reson. Med. 73, 1246–1251 (2015).

60. Shih, Y.-Y. I. et al. Imaging neurovascular function and functional recovery after stroke in the rat striatum using forepaw stimulation. J. Cereb. Blood Flow Metab. Off. J. Int. Soc. Cereb. Blood Flow Metab. 34, 1483–1492 (2014).

61. Lee, S.-H., Ban, W. & Shih, Y.-Y. I. BrkRaw/bruker: BrkRaw v0.3.3. Zenodo 10.5281/ZENODO.3877179 (2020).

